# Moving Beyond Clinical Risk Scores with a Mobile App for the Genomic Risk of Coronary Artery Disease

**DOI:** 10.1101/101519

**Authors:** Evan D. Muse, Nathan E. Wineinger, Brian Schrader, Bhuvan Molparia, Emily G. Spencer, Dale L. Bodian, Ali Torkamani, Eric J. Topol

## Abstract

Primary prevention of coronary artery disease (CAD) is important for individuals at increased risk, and largely consists of healthy lifestyle modifications and initiation of medications when appropriate – including statins. Defining the inherent risk for any given individual typically relies on traditional risk factors established decades ago by the Framingham Heart Study. Unfortunately, recent studies have indicated that these traditional clinical risk factors systematically overestimate the risk of CAD across all major ancestries. This has increased the number of patients that would be eligible for statin therapy for the primary prevention of CAD but would likely receive little benefit and potentially incur negative consequences. On the other hand, researchers have demonstrated that genetic factors can effectively identify a subset of high risk individuals, and that the benefit from statin therapy is greatest among individuals with the highest genetic risk score (GRS). These individuals also receive the greatest absolute benefit from healthy lifestyle choices, being able to titrate their risk to normal levels despite high genetic predisposition. However, it is not yet possible for the average individual to discover their genetic risk because no tools are currently available to make such a determination. Here, we present a free mobile app – MyGeneRank – that can provide this information. Individuals may choose to use this knowledge to complement traditional risk assessments, and make critical decisions regarding lifelong statin therapy and lifestyle changes. As of 1/25/2017, MyGeneRank is currently in closed beta and will soon be available to the public.

## Current Risk Assessment Tools Lack Precision

Clinical determinates relating to risk factors for CAD such as blood pressure, plasma cholesterol and glucose, and cigarette smoking status have been the mainstay in determining a person’s likelihood of developing disease. These factors have been incorporated into tools for tailoring a patient’s individualized prevention strategy but there remains a great deal of uncertainty when applying these clinical risk scores to patient care. In concert with the announcement of guidelines for the treatment of lipid disorders for the prevention of CVD, the American College of Cardiology (ACC) and American Heart Association (AHA) released an online calculator for determining 10-year and lifetime risk of atherosclerotic heart disease on which personalized treatment would be based [1]. Recent studies have indicated that clinical risk factors systematically overestimate the risk of CAD across all major ancestries [2]. In an analysis based on 307,591 adults without type 2 diabetes mellitus, this calculator grossly overestimated the 5-year cardiovascular risk for all groups studied by nearly 5 times that of observed risk [3]. Furthermore, in a head-to-head analysis of five separate ASCVD risk calculators in a multi-ethnic cohort, it was found that 4 of the 5 prediction tools overestimated cardiovascular risk in males (by 37-154%) and females (by 8-67%) for all stratums of risk [4]. When the predicated and observed risk of cardiovascular events for 4,854 participants in the Rotterdam Study was compared using ACC/AHA, ATP-III and ESC criteria, all three independent prediction tools vastly overestimated risk. There are also dramatic differences between the various risk prediction tools and in the expert guidelines from international societies [5]. For example, ACC/AHA guidelines would have recommended statin therapy for 96.4% of men, compared with 52.0% by ATP-III and 66.1% by ESC criteria [6]. Some have argued that overestimation of risk is mainly in higher risk individuals for whom statin therapy would be necessary anyway [7]. Yet better clarity is certainly needed as current risk prediction tools are used daily to decide which patients should initiate lifelong medications despite these gross inaccuracies.

Statins are undisputedly effective in reducing CAD event risk as a secondary prevention strategy [8,9]. But for primary prevention, the net benefit of statins is less clear [10]. Less than 2 out of 100 individuals on statins for 5 years avoid heart attack or stroke, whereas 1 in 100 develop diabetes due to the therapy [11,12]. The choice to go on a lifetime of statins must be individualized. It should not be based simply on guidelines that consider little other than age and gender while severely overestimating risk [3]. In an effort to improve on this, studies have sought to develop genomic risk scores based on variants identified though genome wide association studies (GWAS).

## Clinical Utility of Coronary Artery Disease Genetic Risk Scores

GWAS have identified thousands of genomic variants associated with common diseases and traits [13]. However, the contribution of any individual genetic marker to trait variation across the population remains low [14]. There is still much work to be done to identify all the genetic factors that influence common diseases and we remain unable to accurately and comprehensively predict disease incidence solely based on genetic susceptibility. Due to this limitation and the polygenic nature of most conditions, the genetic risk score (GRS) – an aggregation of estimated genetic risk across multiple loci – has been proposed to translate genomic results into clinical practice. Strong relationships between a GRS and a variety of medical conditions have been found [15], and GRSs have been shown to provide complementary information on disease risk beyond traditional clinical risk factors and are independent of family history [16].

Current estimates of genetic risk in the form of a GRS have been shown to effectively identify high risk individuals and influence behavioral and health decision-making outcomes, particularly related to CAD and the use of statins [17–20]. In one study, the combination of clinical and genetic risk factors double the positive predictive value (PPV) for prediction of incident CAD (43% PPV) vs clinical risk factors alone (23% PPV) [19]. Likewise, other studies showed a reclassification of 12% and 13% of individuals from an intermediate risk category into a high-risk category which would translate into stronger statin-use recommendations and overall reduction in CAD events [17,20]. Thus, knowledge of one’s genetic susceptibility for CAD can help guide the decision to initiate statins, in addition to optimizing overall lifestyle modifications. When presented with their genetic risk for CAD, individuals with higher GRS are more likely to initiate statin therapy [21]. Upon initiation of statin therapy for primary prevention, patients with the highest GRS for CAD achieve the greatest absolute risk reduction of coronary heart disease events with the number needed to treat reduced by more than 2 ½ times as compared to individuals with the lowest GRS [18].

Moreover, the benefits of a patient’s knowledge of their CAD GRS extend beyond the decision to initiate statins. When presented with their genetic risk for CAD, individuals with familial CAD demonstrate higher rates of adherence to therapy [22], and have a higher rate of compliance for performing preventive behaviors [23]. Importantly, patients have the ability to overcome much of their genetic risk for CAD by maintaining optimal lifestyle habits which reduces their risk of disease by nearly half [24]. With such a clear benefit, the challenge instead is in how this information can be relayed to the patient.

## MyGeneRank: A Mobile App for Coronary Artery Disease Genetic Risk

MyGeneRank, a mobile device app which leverages modern data handling processes with a popular genotyping platform and the best practices in genomic sciences, seeks to provide users an unbiased estimate of their genetic risk for CAD to guide in the decision to initiate a statin and encourage efforts of effective lifestyle optimization for primary prevention of CAD. This is the first ever publically available app to deliver this essential personalized data. This no-cost, mobile app may enhance the ability of patients and physicians to make informed medical and lifestyle decisions regarding the primary prevention of CAD. To receive their CAD GRS, users must enroll in an ongoing study to gauge the impact of providing users this information, including behavior and medication changes. Given the rapid pace of discovery in genomic medicine with new genetic markers for disease being identified regularly, users are notified when new disease risk models are developed. However, users have the option to withdraw from these notifications and further study at any time.

MyGeneRank is designed to provide any individual their genetic risk for disease without requiring familiarity in genetic data analysis or data handling. This application interfaces with the 23 andMe application program interface (API) to directly transfer a user’s data to a secure server. A series of computations are then performed on our servers and the user’s estimated genetic risk is returned to them via graphical interface on the smartphone app. GRS is calculated using a weighted allele-counting approach, similar to that performed in prior studies on CAD GRS [18]. Markers and weights were selected based on results from the most current CAD GWAS meta-analysis, which identified 57 markers associated with CAD at the genome-wide level [25]. The markers and corresponding weights are included in **Supplemental Table 1**. Because all of these markers are not typically present on a genotyping array, imputation of these missing markers is performed. The estimated contribution to disease risk from an imputed marker is calculated by treating the genotype probabilities as dosages. The resulting GRS is then compared to a population reference panel, and the resulting percentile rank of genetic risk relative to this population is returned to the user on the app (**Figure 1**).

**Figure 1:**
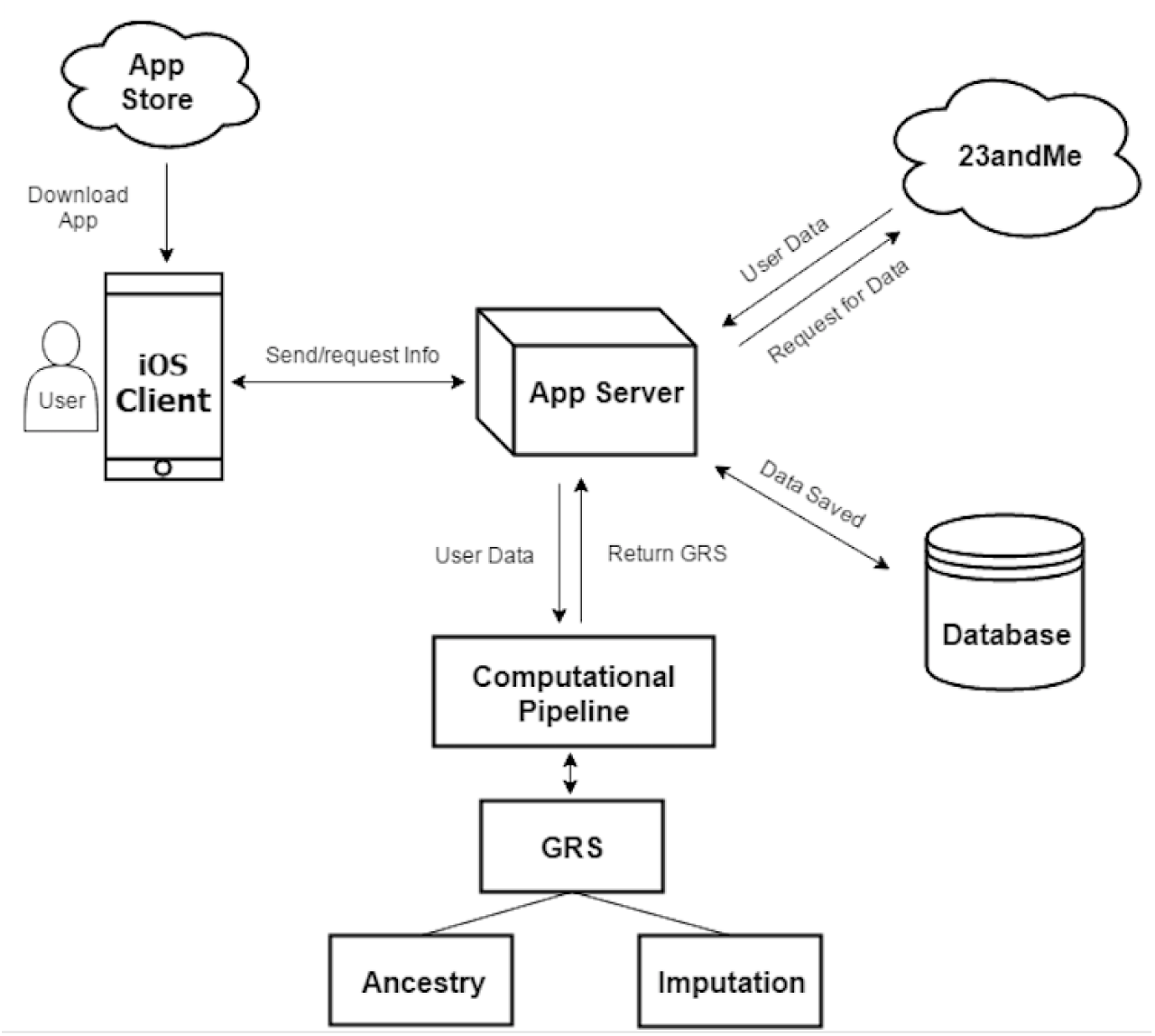
Overview of data acquisition, computation and result delivery

Genetic factors that influence disease have been disproportionately studied in individuals of European ancestry. The most prominent GWAS meta-analyses typically include a large European discovery cohort, but a considerably smaller or non-existent non-European cohort. We discovered that 31% of variation in CAD GRS in 1000 Genomes founders could be explained by continental population alone (**Figure 2**). Considering this, we designed MyGeneRank so that any user regardless of race could learn their unbiased disease risk. First we estimated the ancestral proportions of a user’s genome relative to 1000 Genomes [26]. Then a customized reference panel matching a user’s unique ancestry is simulated from 1000 Genomes, which is used to calculate the user’s genetic risk for disease.

**Figure 2:**
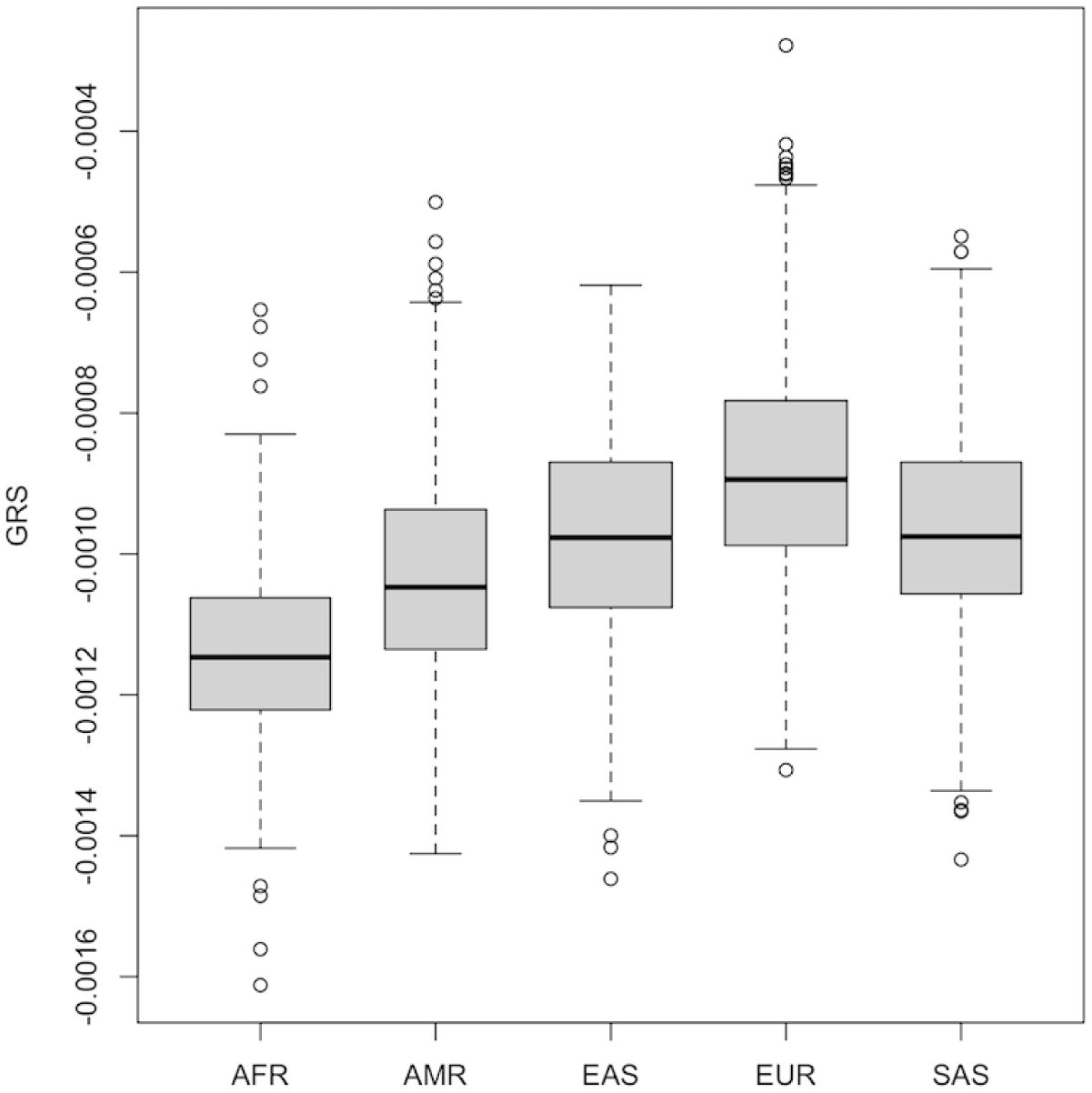
Assessment of CAD GRS in 1000 Genomes cohort based on ethnicity (1000 Genomes super populations). AFR, African; AMR, Ad-mixed American; EAS, East Asian; EUR, European; SAS, South Asian.

To confirm that MyGeneRank provides unbiased estimates of genetic risk, the CAD GRS was calculated using MyGeneRank for 686 population controls as described previously [27]. Briefly, these individuals were healthy volunteers from a pre-term birth cohort, sequenced on Complete Genomics’ sequencing platform, and were 90% or more genetically European. The GRS was computed for each individual, and those scores were compared to scores from the 1000 Genomes reference panel to calculate percentile ranks (**Supplemental Figure 1**). These scores did not statistically differ from the CEU (Caucasians from Utah) reference panel in 1000 Genomes (p=0.35) or all individuals of European ancestry (p=0.08). The mean and median percentiles of estimated CAD genetic risk were 47% and 46%, respectively. These results confirm that MyGeneRank provides unbiased estimates of genetic risk in the population.

The primary goal of MyGeneRank is not the development of novel methods for estimating genetic risk, but is instead to empower patients and physicians with a tool that utilizes previously reported findings to provide information that can be applied in health decision making. Using a smartphone, the app can be accessed almost anywhere, including most outpatient clinics during a physician visit (**Figure 3**). Though the decision to begin statin therapy should be weighed carefully, it was imperative to us that individuals could request and gain access to their CAD GRS with rapid return of results. Therefore, much of the data-handling and computational processes were designed to complete as quickly and efficiently as possible resulting in a turn-around time of approximately 4 minutes. Multiple requests can be handled simultaneously, and a queuing priority system is in place for when the number of requests exceeds the computational load.

**Figure 3:**
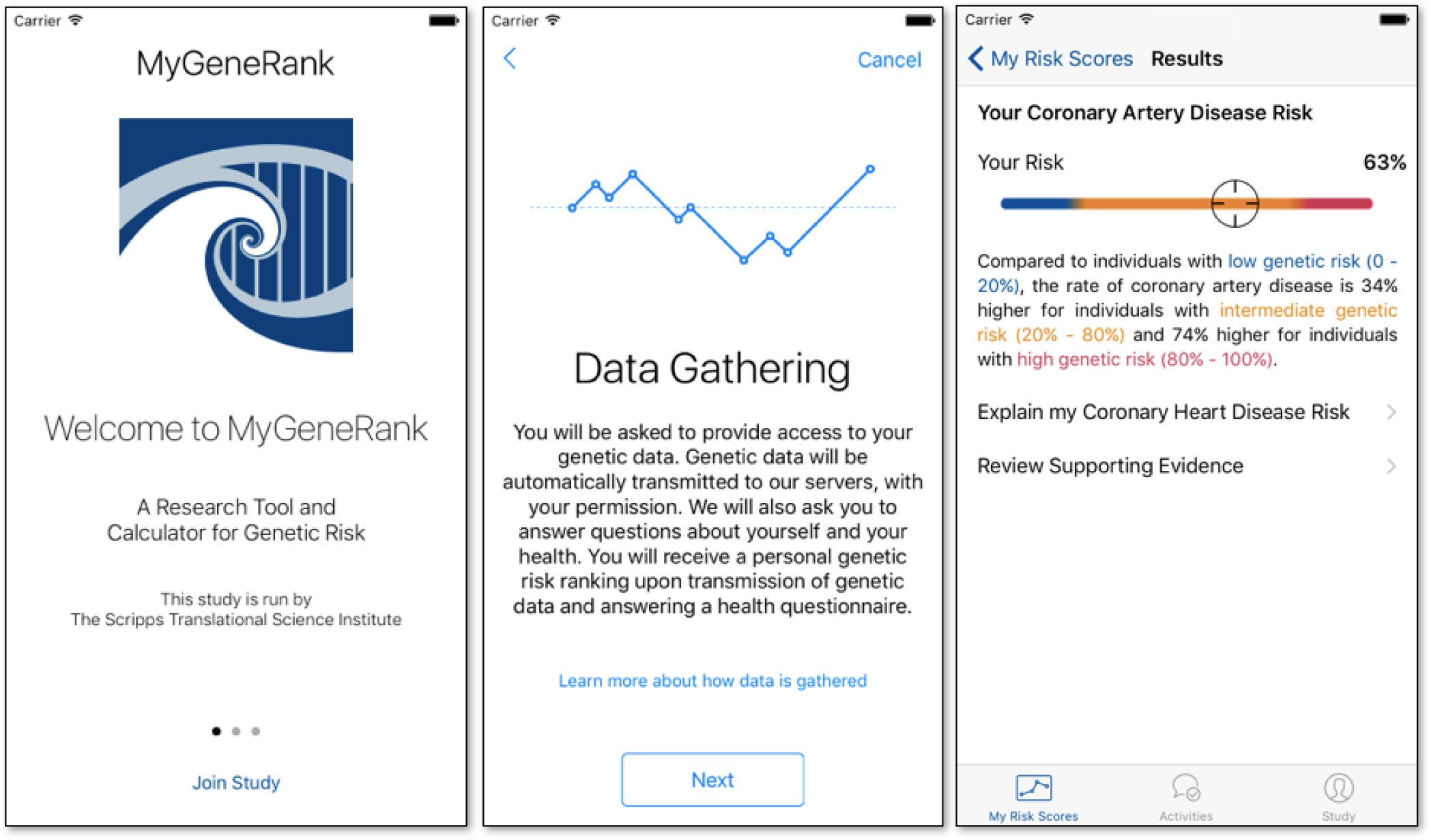
Screen shot from MyGeneRank App

Undoubtedly, new discoveries will emerge and we expect to regularly update the GRS software to reflect our best understanding of genetic risk. As of 1/25/2017, MyGeneRank is currently only available on Apple iOS, but will be updated in the future to be accessible from the web and Android smartphones.

## Incorporating Genetic Risk into Overall Cardiovascular Risk Assessment

Despite rapid advances in genomic medicine that are providing novel insights into the pathogenesis of rare and common diseases, integration of this data into everyday clinical practice has been slow. With the continuing debate over which individuals should and should not be taking statins for primary prevention of cardiovascular disease [28] and the potential benefits of using a GRS, it is an appropriate time to introduce a CAD GRS in the health decision-making process. It is clear that currently used clinical risk calculators for CAD are inadequate, especially as it relates to starting a medication that may last a lifetime [3] or vital information that would potentially alter one’s lifestyle, including nutrition and physical activity. Many patients seeking to fully optimize their CAD primary prevention strategies work to balance their risk of developing disease with the implications of long-term or even life-long statin therapy or steadfastness with aerobic activity and healthy diets. Unfortunately, based on clinical risk factors alone, many people fall into a gray zone where the data falls short and many individuals opt to avoid statin therapy given the numerous reports of adverse side effects in the press. For an otherwise healthy patient solely with borderline-elevated cholesterol measurements, the CAD GRS would certainly serve as a powerful determining factor in the overall risk-benefit discussion. For patients with the highest genetic risk for CAD (and as previously discussed the lowest NNT on statin therapy) the scales would likely not only tip in favor of initiation of pharmacotherapy, but those patients would have higher compliance rates.

As GRSs have been derived for multiple disease models beyond CAD there is potential to expand to other diseases as well. Though the current version of MyGeneRank requires users to provide genetic data through a 23andMe account, future versions will allow for other sources of genotype and sequencing data. Not only will individuals be armed with their own data in a way that is easily accessible and interpretable from almost anywhere, but participants will have the opportunity to seamlessly participate in a real-world validation study that may complement the way we approach cardiovascular disease prevention moving forward. By simply downloading the app and consenting to sharing of their self-reported disease status and genetic data with personal identifiers removed, volunteers will be participating in a new generation of clinical trials driven by and for patients. Such a capability may help translate a large body of vital knowledge that sits in the research domain to the real world to help individuals make decisions about their lifestyle and medications.

## KEY MESSAGES

- Which patients should be treated with statins for the primary prevention of cardiovascular disease remains debated.
- Genetic risk scores for coronary artery disease have been shown to identify patients at increased risk of CAD and who would achieve the greatest benefit of statin therapy.
- Despite this, there are no realistic ways for the average individuals to learn their genetic risk
- MyGeneRank is a free, mobile app that provides this information and can be used to inform the decision-making process in regards to statin therapy and lifestyle optimization.

## CONTRIBUTORS

EDM, NEW, AT, and EJT conceived of the conceptual framework. NEW, BS, EGS, and AT designed the app. BS developed the app. NEW and BM developed the GRS pipeline. DLB provided population controls. AT and EJT obtained funding. EDM drafted the manuscript and NEW, AT, and EJT provided editing. All authors approved the manuscript for publication.

---

**Supplementary Figure 1:**
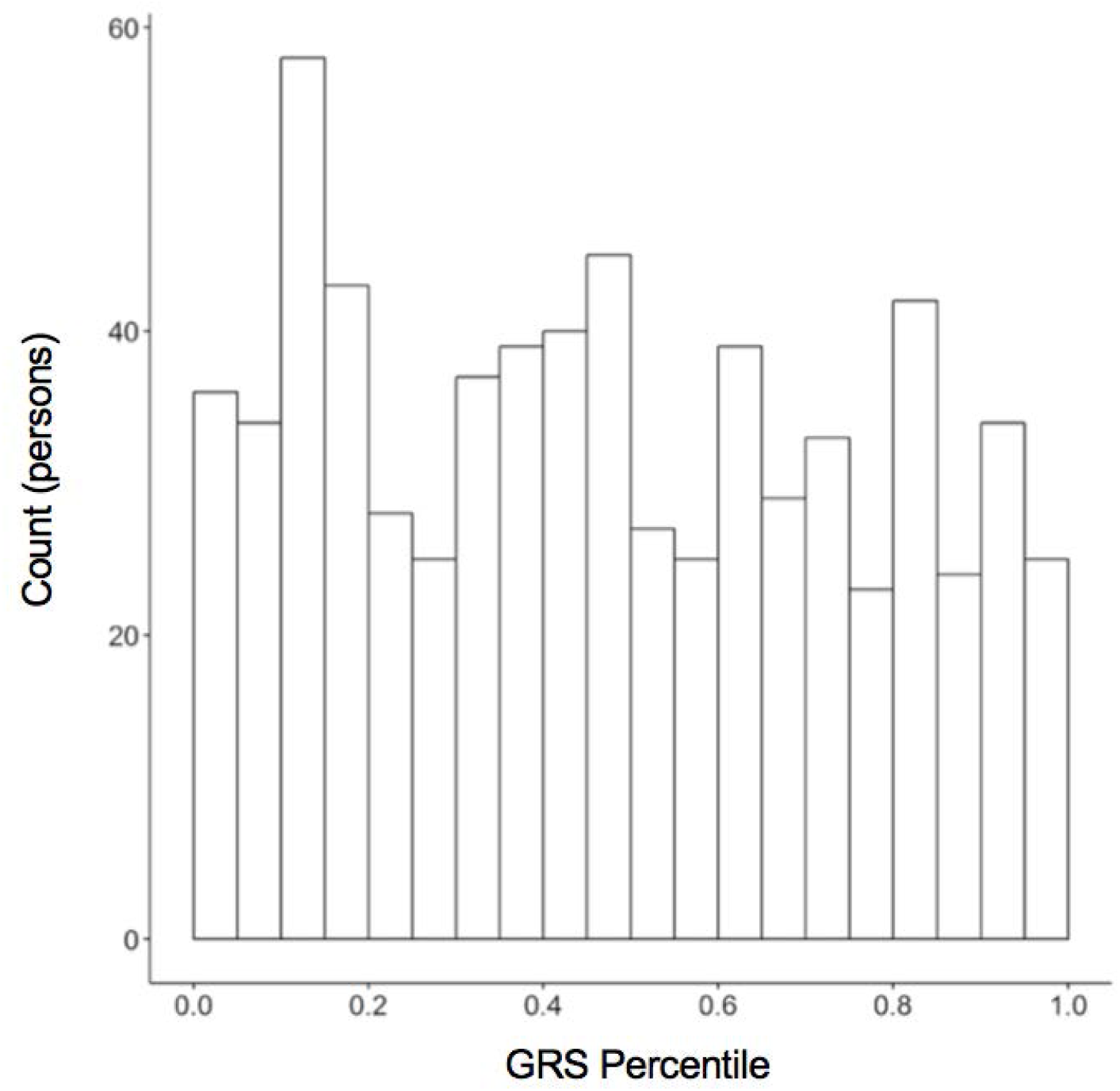
Screen shot from MyGeneRank App

